# Allocation of visuospatial attention indexes evidence accumulation for reach decisions

**DOI:** 10.1101/2022.05.06.490925

**Authors:** Carolin Schonard, Tobias Heed, Christian Seegelke

## Abstract

Visuospatial attention is a prerequisite for the performance of visually guided movements: Perceptual discrimination is regularly enhanced at target locations prior to movement initiation. It is known that this attentional prioritization evolves over the time of movement preparation; however, it is not clear whether this build-up simply reflects a time requirement of attention formation or whether, instead, attention build-up reflects the emergence of the movement decision. To address this question, we combined behavioral experiments, psychophysics, and computational decision-making models to characterize the time course of attention build-up during motor preparation. Participants (n = 46, 29 female) executed center-out reaches to one of two potential target locations and reported the identity of a visual discrimination target that occurred concurrently at one of various time-points during movement preparation and execution. Visual discrimination increased simultaneously at the two potential target locations but was modulated by the experiment-wide probability that a given location would become the final goal. Attention increased further for the location that was then designated as the final goal location, with a time course closely related to movement initiation. A sequential sampling model of decision-making predicted key temporal characteristics of attentional allocation. Together, these findings provide evidence that visuospatial attentional prioritization during motor preparation does not simply reflect that a spatial location has been selected as movement goal, but rather indexes the time-extended, cumulative decision that leads to selection, hence constituting a link between perceptual and motor aspects of sensorimotor decisions.

**Significance statement:** When humans perform a goal-directed movement such as a reach, attention shifts towards the goal location already before movement initiation, indicating that motor goal selection relies on the use of attention. Here, we demonstrate that key temporal aspects of visuospatial attention are predicted by a well-known computational model of decision-making. These findings suggest that visual attention signals much more than simply that a motor goal has been selected: instead, the time-course of emergent, visuospatial attention reflects the time-extended, cumulative decision that leads to goal selection, offering a window onto the tight link of perceptual and motor aspects in sensorimotor decision-making.

## Introduction

Our environment usually presents us with multiple, concurrent action opportunities. Successful interaction, therefore, continuously requires decisions about motor goals (i.e., what to do) and the specification of the respective motor parameters (i.e., how to do it; Cisek and Kalaska, 2010; Wolpert and Landy, 2012; Wong et al., 2015; Scott, 2016; Gallivan et al., 2018).

The selection of motor goals relies on (visuospatial) attention (Allport, 1987; Awh et al., 2006; Wong et al., 2015; Li et al., 2021), evident in improved perceptual discrimination performance at movement target locations compared to other locations. Attention shifts towards movement targets already prior to movement initiation and reliably occurs during the preparation of saccadic eye movements (Hoffman and Subramaniam, 1995; Kowler et al., 1995; Deubel and Schneider, 1996; Collins et al., 2010; Rolfs and Carrasco, 2012) and reaching movements (Deubel et al., 1998; Baldauf et al., 2006; Collins et al., 2008). Neurophysiological and neuroimaging evidence corroborate these attention-related behavioral improvements by showing modulation of neural activity in (oculo)motor-related brain structures, such as the frontal eye fields (FEF), the superior colliculi (SC), and the lateral intraparietal areas (LIP), during visuomotor attention tasks (Corbetta et al., 1998; Nobre et al., 2000; Moore and Fallah, 2004; Bisley and Goldberg, 2010; Wardak et al., 2011; Bollimunta et al., 2018), further underscoring the tight coupling between motor preparation and spatial attention.

Remarkably, participants can divide their attention between multiple target locations simultaneously (Baldauf et al., 2006; Baldauf and Deubel, 2008; Hanning et al., 2018). These findings fit the concept of a dynamic attentional landscape (Baldauf and Deubel, 2010) or priority map (Wolfe, 1994; Fecteau and Munoz, 2006) that is continually constructed through bottom-up and top-down inputs. Activity within this map represents a spatial layout of available options, weighted by their behavioral relevance. Activity peaks allow for selection of motor goals and guidance of visual attention on a moment-by-moment basis (Bisley and Goldberg, 2010). Critically, this framework suggests that visuospatial attention is not the result (and, thus, a marker) of a finalized selection process, but, instead, constitutes a link between perceptual and motor aspects of sensorimotor decisions that lead to selection of these goals.

In line with this proposal, a recent study demonstrated that attentional allocation emerged continuously during motor goal selection (Jonikaitis et al., 2017). Participants prepared saccades to two potential targets, of which one was cued as the final target only after a delay. Visual attention, probed via the sensitivity for visual discrimination, was elevated at both pre-cued locations, and gradually increased at the final target, temporally linked to saccade onset. The authors proposed that the spatiotemporal properties of attentional allocation reflected oculomotor decision-making as put forward in sequential sampling models, in which evidence gradually accumulates over time until it reaches a threshold that elicits an overt response (c.f. Gold and Shadlen, 2007; Ratcliff et al., 2016).

Here, we provide a rigorous test of this assertion. We first extend the delayed-cueing paradigm to hand reaches and demonstrate that the gradual increase of visuospatial attention and its distribution across relevant locations is a general computational principle across effector systems. We then show that the probability with which a given target will later become the instructed movement goal – a variable known to affect decision-making (Platt and Glimcher, 1999; Ratcliff et al., 1999; Hudson et al., 2007; Wong et al., 2022) – modulates both the spatiotemporal characteristics of attentional allocation and reach behavior, indicating that top-down information about upcoming actions alters attentional prioritization. Finally, we bridge attentional and motor aspects by fitting a sequential sampling model of decision-making to the movement data. The derived predictions account for key temporal aspects of attentional allocation. Together, our results provide compelling evidence that in situations of target uncertainty, visuospatial attention during motor preparation does not just reflect ultimately selected motor goals, but rather the time-extended, cumulative decision leading to their selection.

## Methods

### Overview of task and participant selection

The three main experiments required participants to report the identity of a visual discrimination target that was presented during a reaching task. Discrimination performance served as a marker for visuospatial attention (e.g., Baldauf et al., 2006; Collins et al., 2008). We adjusted the duration of the discrimination target between 50 and 200 ms to each participant’s perceptual thresholds, determined in a separate session prior to the main experiment. We excluded participants from the main experiments if they did not reach 85% accuracy with the longest allowed stimulus duration of 200 ms.

### Participants

Previous studies with similar paradigms yielded consistent results with approximately 10 participants (e.g., Baldauf et al., 2006; Baldauf and Deubel, 2008; Jonikaitis et al., 2017; Hanning et al., 2019; Wollenberg et al., 2020). We thus decided to obtain data from 15 participants for each of our three experiments. We first screened participants’ ability to reliably perform the perceptual discrimination task employed to assess attention. We screened 123 physically and neurologically healthy individuals from Bielefeld University (Experiment 1: 47 participants, Experiment 2: 48 participants, Experiment 3: 28 participants). 48 (Exp. 1: 21; Exp. 2: 17; Exp. 3: 10) participants did not meet the required visual discrimination performance criterion (see below) and thus were not further tested. Another 15 (Exp. 1 6; Exp. 2 6; Exp. 3 3) participants made excessive eye movements and were removed from our samples after participation in the experimental sessions. Yet another 15 (Exp. 1 4; Exp. 2 9; Exp. 3 2) participants did not finish the experiment. The final samples consisted of 15 participants for Experiment 1 (9 female, mean age = 22.4 years, SD = 3.1 years; 14 right-handed, mean handedness score = 99.12; 1 left-handed, mean handedness score = -62.5; Dragovic, 2004), 16 participants for Experiment 2 (7 female, mean age 23.1 years SD = 3.5 years; 15 right-handed, mean handedness score = 89.51; 1 left-handed, mean handedness score = -42.86), and 15 participants for Experiment 3 (13 female, mean age = 21.3 years SD = 2.4 years; all right-handed, mean handedness score = 100). The ethics committee at Bielefeld University approved the experiments (Ethical Application Ref: 2018-155). All participants gave their informed written consent to participate in the study and were naïve to the purpose of the experiment. They received course credit or 7 € per hour in exchange for their participation.

### Apparatus, stimuli, and task

Participants sat on a chair and grasped the handle of a robotic manipulandum (KINARM End-Point Lab, BKIN Technologies, Kingston, ON, Canada) to perform center-out reaching movements in the horizontal plane. They controlled the handle with their right hand to move a circular white cursor (diameter = 1.0 cm) from a starting position to “shoot” through a target. Stimuli and cursor were projected from a horizontal screen with a 60 Hz refresh rate onto an opaque mirror. Note that the screen refresh rate limited all visual stimulus durations to multiples of 16.6 ms. The mirror was placed halfway between the screen and the arm such that the stimuli and cursor appeared on the same plane as the handle, but the participant’s arm was hidden from view by the mirror. The KINARM recorded handle position, velocity, and acceleration at 1000 Hz.

#### Main task

##### Experiment 1

Each trial started with the presentation of the stimulus display. This display contained a central circular grey starting position with 4.0 cm diameter that was located approximately 30 cm vertically from the middle of the participant’s chest and twelve pre-masked characters resembling a digital number ‘8’ (3 cm in height, 2 cm in width; see Figure 1A). The characters were arranged equidistantly on an imagery circle of 10.0 cm radius around the starting position. After participants had maintained the cursor at the starting position for 1,500 ms, two 4 cm diameter, equiluminant colored circles pre-cued two character locations for 500 ms. One circle was blue and the other green. These pre-cued locations indicated the two possible reach targets of the current trial. Reach targets could appear at 1, 3, 5, 7, 9, and 11 o’clock (cf. Baldauf et al., 2006). The pre-cues were followed by a delay period of 500 ms to 1000 ms that was randomly drawn from a uniform distribution; the color pre-cues were extinguished during this time. After the delay period, the central starting position was illuminated with the color of one of the two pre-cues for 500 ms; both color cues occurred equiprobably. This cue specified the reach target, and participants had to “shoot” the cursor through the respective target location as fast as possible. Concurrently, a probe display was presented at a variable onset time between the onset of the delay period and 500 ms after central color cue onset. This time point was randomly drawn from a uniform distribution with a step size of 16.6 ms. Individual probe display duration ranged from 50 to 200 ms (Experiment 1 median = 167 ms, Experiment 2 median = 133 ms, Experiment 3 median = 117 ms), adjusted to the individual participants’ performances in the threshold task (see below). In the probe display, eleven of the twelve pre-mask target characters took on the shape of a digital number ‘2’ or ‘5’, and one character, the discrimination target (DT), took on the shape of a digital number ‘3’ or the letter ‘E’. After the probe presentation time had passed, all characters changed back to the digital ‘8’ mask. At the end of each trial, participants reported whether the DT was a ‘3’ or ‘E’ by pressing one of two response buttons (Buddy Button, Ablenet, Roseville, MN, USA) with their left hand. The DT could appear at any of the six target positions with equal probability. Thus, in relation to the movement task, the DT could appear at the finally cued, *instructed* target location, at the other pre-cued, but *discarded* target location, or at one of the four non-cued locations that were *irrelevant* for movement in the present trial. We used an electrooculogram (EOG) to ascertain that participants maintained fixation at the central starting position throughout the entire trial (see below).

**Figure 1.**
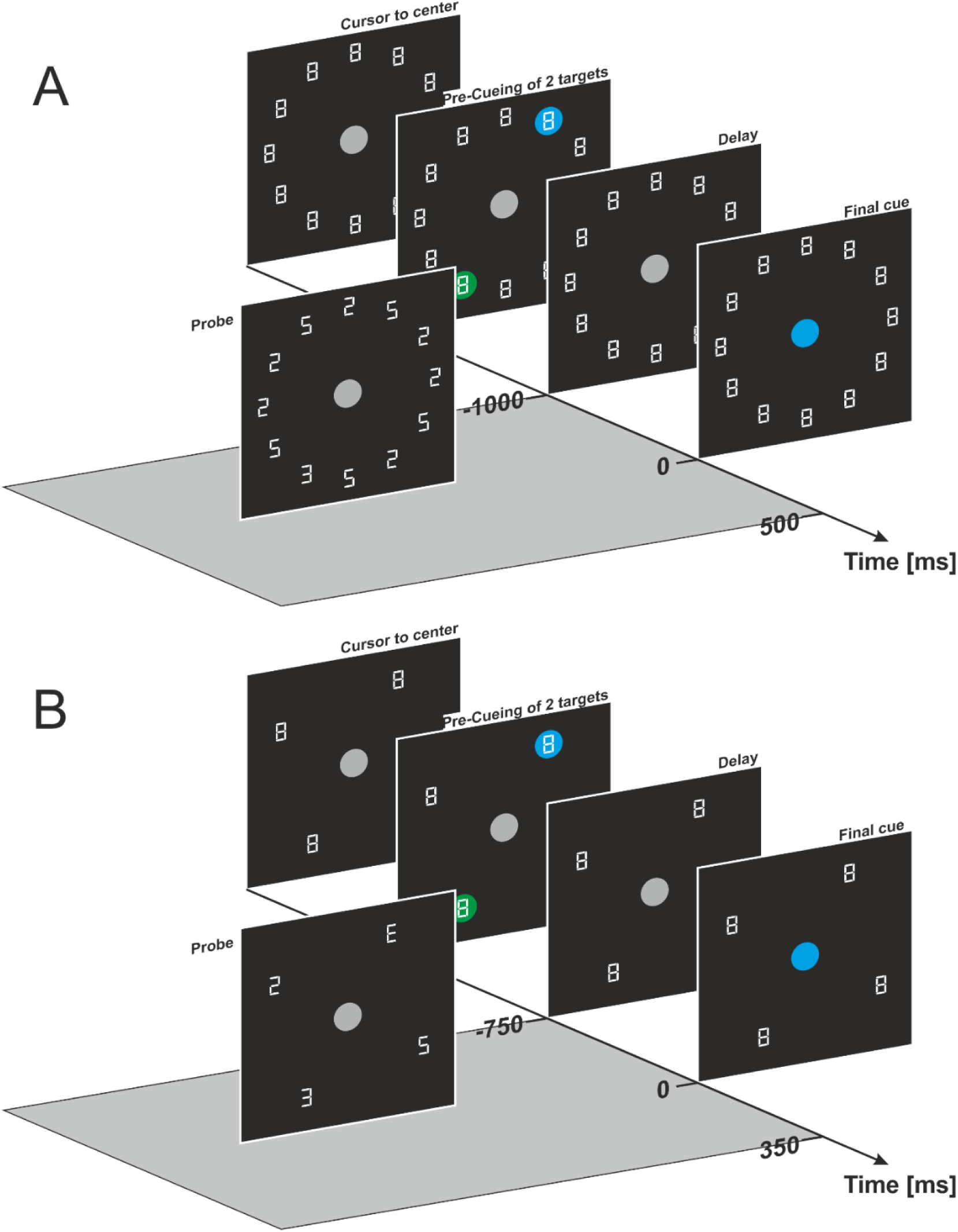
Stimulus display and trial structure of Experiments 1 and 2 (**A**), and Experiment 3 (**B**). Upon moving the robotic device into the grey center circle, two reach target locations were marked in blue and green color. After a variable delay, the central circle changed to one of the two colors and participants had to reach towards the location that had previously been marked with this color. Concurrently, the probe display replaced the digital ‘8’s with ‘5’ and ‘2’ distractors and one ‘E’ or ‘3’ discrimination target. In Experiment 3, only four target locations were used, and two discrimination targets (instead of one) were displayed for the matching task.

##### Experiment 2

Stimulus presentation was identical to that of Experiment 1 with the following exceptions: Based on the results of Experiment 1, we shortened the duration of the delay period to 500-750 ms, and the probe display was presented at a variable time between the onset of the delay phase and 300 ms after final cue onset. The main manipulation of Experiment 2 was that the two pre-cued targets turned into the final reach target with different probabilities. One pre-cue color (e.g. blue; balanced across participants) indicated that the respective target location was finally instructed as reach target with a probability of 80%; accordingly, the other pre-cue color (e.g., green) indicated that the respective location was instructed as reach target with 20% probability. Participants were not informed about the probability manipulation but could learn these probabilities through experience.

##### Experiment 3

In Experiment 3 we replaced the discrimination task of Experiments 1 and 2 with a matching task (Figure 1B). Stimulus display and task procedure were identical to Experiment 2 with the following exceptions: Only four characters, rather than 12, were arranged on the imagery clock positions of 1, 4, 7, and 10 o’clock around the central starting point. All four locations could be selected as reach targets. In the probe display, two of the four pre-mask characters took on the shape of a digital number ‘2’ or ‘5’, and two pre-mask characters (the DTs) took on the shape of a digital number ‘3’ or ‘E’. At the end of each trial, participants reported whether the DTs were identical or different by pressing one of two response buttons with their left hand.

In all experiments, participants performed between 5400 and 5580 trials, split into blocks of 180 trials. Within each block, each target location was repeated 30 times (Experiment 1 and 2) or 45 times (Experiment 3), presented in a randomized order. Data acquisition was spread across five to six sessions, and participants performed between three and seven blocks per day with each session lasting about 2.5 hours.

#### Threshold task

Visual discrimination performance can differ considerably across participants (Baldauf et al., 2006; Hanning et al., 2019). Therefore, we determined the probe display duration for each individual on a separate day prior to the main experiment. To this end, we assessed at which presentation duration the participant could discern the discrimination target with 85% accuracy. Stimulus presentation was identical to the main task. However, the discrimination target always appeared at the pre-cued location, and only a single target was pre-cued. Participants were explicitly informed about this contingency. The duration of the probe display varied randomly sampled from a uniform distribution between 50-350 ms (participants 1-22) and 50-300 ms (from participant 23 on), respectively, in steps of 16.6 ms. Participants performed one practice block that was not analyzed; they then performed 4 or 5 blocks, with each block comprising 180 trials. For the main task, we then employed the probe duration at which a participant just exceeded 85% correct discrimination performance (Baldauf et al., 2006). Note that perfect adjustment of probe presentation times to 85% discrimination accuracy was not possible because the monitor refresh rate restricted visual stimulus durations to multiples of 16.6 ms.

#### EOG recording

The EOG was recorded from Ag/AgCl electrodes placed above and below the right eye (EOG_up_ and EOG_down_) and at the outer canthi of the left and right eye (EOG_left_ and EOG_right_). The electrodes were referenced to the right mastoid and grounded to the forehead. The EOG signal was recorded with a BrainAmp DC amplifier/ BrainAmp MR DC amplifier (Brain Products GmbH, Gilching, Germany) and digitally stored using the BrainVision Recorder software (Brain Products GmbH). The analog EEG signal was sampled at 5000 Hz, filtered online with a bandpass of 0.1 to 250 Hz, and then down-sampled online to 500 Hz.

#### Acquisition of EOG training data

Participants performed a short saccade task at the beginning of each recording session to obtain training data for a probabilistic classifier that detects blinks and saccades (Toivanen et al., 2015). Stimulus presentation was similar to the main and threshold task, but no movement with the robotic manipulandum was required. At the start of each trial, participants fixated the central starting position. After 1,500 ms, one of the six target positions was illuminated in blue or green color for 2,000 ms, and participants made a saccade towards the respective location and maintained fixation there. Then, the starting position was illuminated, and participants made another saccade and fixated that position. After 2,000 ms, the colored circle was extinguished, and following a 2,000 ms inter-trial interval, the starting position became grey to indicate the start of the next trial. Participants performed 5 saccades to each of the six possible target positions presented in randomized order.

#### EOG analysis

Blinks and saccades were classified with a probabilistic algorithm that calculates the probability that a given sample contains a fixation, saccade, or blink (Toivanen et al., 2015). In this approach, an expectation-maximization algorithm is used to determine the parameters of a Gaussian model in an unsupervised learning phase. If the probability mass for blink or saccade samples exceeded 90%, we marked the blink or saccade as detected.

### Data processing and analysis

We filtered kinematic data of the reaching movements using a 3rd-order zero-lag double-pass filter with a cutoff frequency of 10Hz. We determined reach onset as the time of the sample in which the velocity exceeded 20 mm/s, and reach offset as the time of the sample in which the cursor entered the target location. We defined reaction time (RT) as the time between final cue and reach onset, and movement time (MT) as the time between reach on- and offset.

We removed trials from analysis in which participants initiated their movements before final cue onset, did not hit a target within 2,000 ms, or made a saccade or a blink between pre-cue onset and DT offset. In addition, we excluded trials when RTs or MTs deviated more than 2.5 SDs from the design cell mean, and trials with RTs above 1,000 ms or MTs above 500 ms. Overall, we excluded 15.0%, 17.5%, and 14.7% of trials for Experiment 1, Experiment 2; and Experiment 3, respectively.

#### Statistical approach

We used response accuracy in the discrimination task as a marker for the allocation of visuospatial attention. Response accuracy was defined as the percentage of correctly identified discrimination targets within a given experimental condition. Chance level was at 50% because participants chose between two response alternatives (‘E’ or ‘3’, and ‘same’ or ‘different’). The time point of DT presentation was defined as the midpoint of its presentation interval. We used the ‘SMART’ method (van Leeuwen et al., 2019) to construct the time course of discrimination accuracy in relation to (1) final cue onset and (2) reach onset, separately for each level of the factor DT location and collapsed across all movement target locations. To this end, pairs of time and discrimination performance data across all trials per participant served as input to a moving Gaussian kernel (σ = 30; step size 1 ms), resulting in a smoothed and continuous time course per participant that reflects response accuracy, that is, the proportion of correct discrimination, across the time of a trial. The individual time-course data were then averaged across participants using a weighted mean, assigning more weight to participants who contributed a larger amount of data at a given time point. We determined 95% confidence intervals for the accuracy differences between the DT locations for each pairwise comparison and calculated *t*-tests at each point in time. If *t*-tests were significant for more than two consecutive time steps, they were marked as a cluster, and the cluster strength was determined by the sum of *t*-values within the cluster. Next, we obtained 1,000 permutations of our data by shuffling the labels that assign the data to experimental conditions. For each permutation, we applied the same analysis steps as for the non-permuted data: Gaussian smoothing and averaging, identifying clusters with significant differences, and calculating the cluster strengths. A permutation distribution was then built by using the cluster with the highest strength from each permutation. If there were no significant clusters within a permutation, the largest *t*-value was used instead. The *p*-value of the clusters obtained in our original, non-permuted data is given by the proportion of permuted clusters with equal or higher cluster strengths.

### Modeling

We modeled the sensorimotor decision process that produced the observed reach behavior in Experiments 1 and 2 within the framework of the leaky competing accumulator (LCA) model (Usher and McClelland, 2001) by fitting the model to reach latency (i.e., RT) and accuracy data. The general idea of accumulator models is that decisions are made by accumulating and integrating evidence in favor of each potential response in latent accumulators which are dedicated to each response option. In the LCA model, evidence is thought to deteriorate, or “leak” with time; this feature discounts temporarily confined in favor of continuous, longer-term evidence. Moreover, the different accumulators inhibit each other in proportion to their accumulation. When the evidence collected in one accumulator, termed “activation”, crosses a predefined threshold, the associated response is selected, marking the decision for the respective option. In support of this idea, neural activity in the posterior parietal cortex exhibits ramping activity that seems to reflect evidence accumulation during perceptual decision-making (c.f. Gold and Shadlen, 2007).

In the LCA model, the activation of the *i*th accumulator is represented by the equation

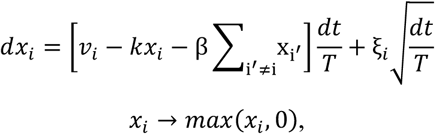

where *v*_*i*_ is the accumulation rate for the *i*th accumulator; ξ_*i*_ is a Gaussian noise term with a mean of zero and standard deviation σ^2^; β is the lateral inhibition exerted on all other accumulators; *k* is the leakage rate of information during the evolving decision; and *T* is the time scale required for integration. We set the time scale to 0.001, indicating 1 ms time steps. We fitted the model with a maximum likelihood approach, using the Probability Density Approximation (PDA) method (Turner and Sederberg, 2014). In short, a synthetic dataset is simulated, and the simulated observations are used to construct the likelihood approximation using a kernel density estimate (for details, see Holmes, 2015). We maximized the likelihood by first performing a grid search within plausible parameter ranges (see appendix), where we evaluated about 12,000 parameter combinations. The 10 best parameter combinations were further optimized with a simplex algorithm, from which we selected the parameter combination that resulted in the highest likelihood.

We included two accumulators: One for (correct) responses to the instructed reach target, and the other for (incorrect) responses to the discarded reach target while excluding incorrect reaches to non-pre-cued locations from fitting (0.3% for Experiment 1, 0.2% for Experiment 2). We expected the time course of the model’s accumulation to be similar to the time course of perceptual discrimination performance. For scaling the model, we let the accumulation rate for the correct response (*v*_*correct*_) vary freely between 0 and 1. *v*_*incorrect*_ was defined as 1 − *v*_*correct*_. The parameters for leakage, inhibition, and noise varied freely, but were shared between the two accumulators. To determine the decision, we added the response threshold *α*, which had to be exceeded by one of the accumulators within a maximum decision time of 1.5 s, and a parameter for the non-decision time that reflects non-decisional perceptual and motor processes.

## Results

### Experiment 1

We examined whether the temporal dynamics of covert visual attention reflect decisional processes of motor goal selection. To this end, we analyzed the time-course of perceptual discrimination performance in response to briefly flashed discrimination targets (DTs) at a pre-cued and consecutively *instructed* reach target, another pre-cued but later *discarded* target, and movement *irrelevant* locations.

Figure 2A shows the time course of response accuracy in the discrimination task time-locked to the onset of the final cue. First, we examined whether perceptual discrimination performance at the two pre-cued, potential reach target locations was enhanced during the delay period, that is, when the final reach goal was still uncertain. Compared to irrelevant locations, discrimination performance was better both at the pre-cued and then instructed reach target (cluster-based permutation test: *p* < .001) and at other pre-cued but then discarded target (*p* < .001). Thus, initially, both pre-cued target locations were attentionally selected in parallel and hence considered as movement targets. Inspection of Figure 2A suggests that discrimination performance was better at the later instructed than at the discarded target location even before presentation of the final cue. While the sequential t-tests in this time window reached significance (all p < .05), analysis of the cluster did not (p = .112). At first glance, this finding is surprising because participants did not know which target would be reach-relevant at this time. However, others have reported similar observations and attributed them to reactivation of working memory contents by post-cued attention (Hanning et al., 2019). In particular, cueing attention towards a stimulus location as late as 900 ms after flashing a stimulus there can improve an observer’s capability to perceive its presence and orientation (Astle et al., 2012; Sergent et al., 2013). It is, thus, conceivable that discrimination performance was affected by such attentional and working memory mechanisms.

**Figure 2.**
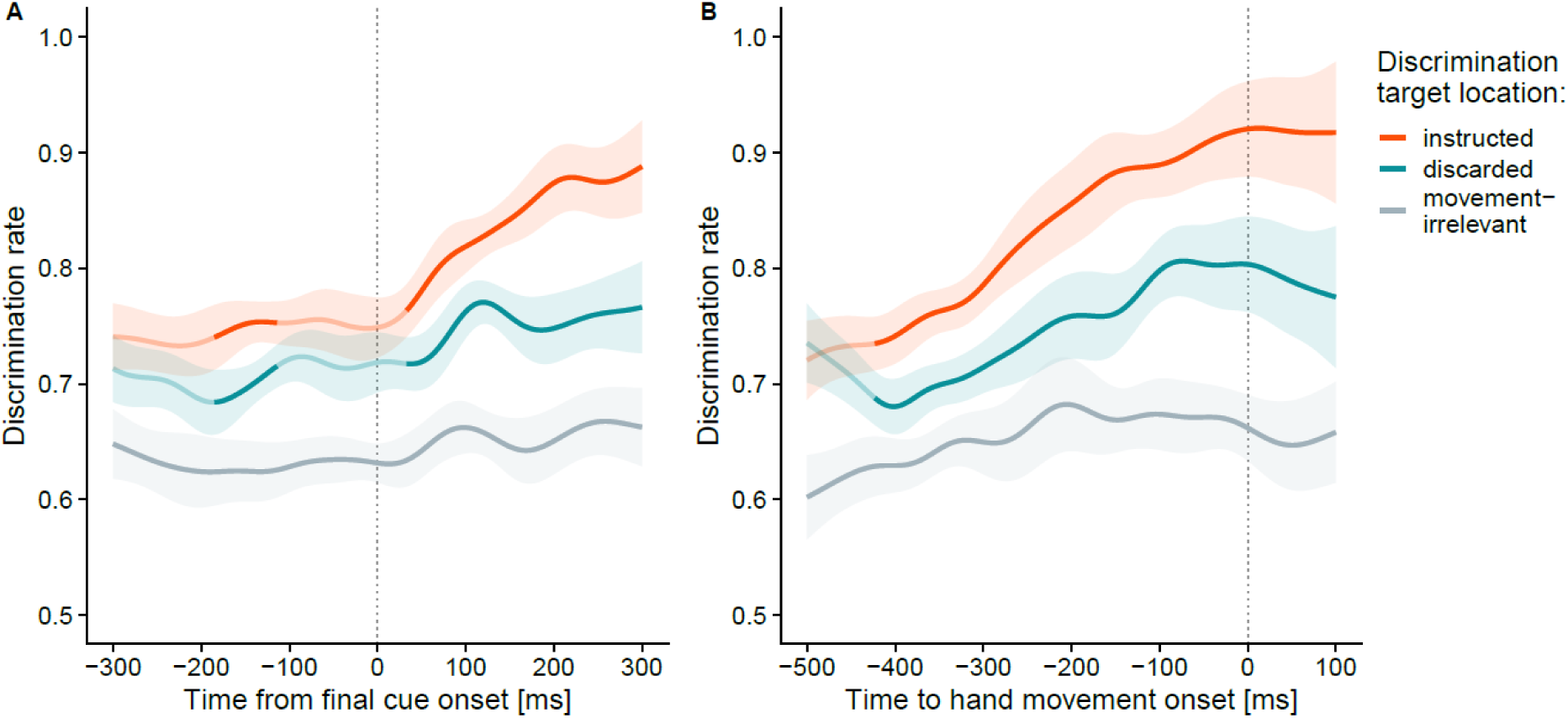
Time-course of visuospatial attentional in Experiment 1. Time-smoothed discrimination rates are shown separately for trials in which the DT appeared at the instructed movement target location (red), at the discarded target location (blue), and at movement irrelevant locations (grey), time-locked to final cue onset (**A**) and reach onset (**B**). Data are smoothed and weighted by the number of available responses at each time point across participants (see Method section). Saturated segments denote time periods with significant differences (*p* < .05, time-point-wise *t*-tests) between instructed and discarded target locations (red/blue) and between instructed target and movement irrelevant locations (grey), respectively. Shaded areas indicate the 95% confidence intervals for the respective pairwise comparisons.

Next, if perceptual discrimination performance at target locations reflects decisional processes related to the selection of a motor goal, we should expect a gradual increase of perceptual performance after designation of the final target. In line with this reasoning, shortly after onset of the final cue discrimination performance increased monotonically at the instructed target and was superior to the discarded target (at ∼35 ms; *p* < 0.001). Somewhat surprisingly, perceptual performance also slightly increased at the discarded target location after final target designation. One might have expected that it would no longer be attentionally selected, given that this location was no longer a potential reach goal. We will return to this point in the interim summary.

Next, we reasoned that discrimination performance should be related to overt reach behavior. To test this conjecture, we analyzed discrimination accuracy time-locked to (participant-determined) reach onset rather than (experimentally dictated) final cue presentation. Discrimination performance rose gradually and monotonically for both instructed and discarded target locations and peaked at reach onset (Figure 2B). Performance was superior at the instructed compared to both the discarded target (*p* < .001) and movement irrelevant locations (*p* < .001), again underscoring the preferential processing at movement goal locations.

Visual inspection of Figure 2B suggests that, on average, performance was highest at the time of reach onset, revealing a correlate of committing to the decision. If this reasoning is correct, we should also expect that reaches initiated at lower latencies would be preceded by either a steeper or earlier rise of attentional allocation (or a combination of the two) compared to reaches initiated at higher latencies, because evidence will be accumulated faster or earlier (or both) prior to fast decisions. Accordingly, we sorted trials into “fast” and “slow” trials at the median RT separately for each participant. Mean RT was 299 ms (SD = 73 ms) in the fastest 49.6% of trials and 465 ms (SD = 132 ms) in the slowest 50.4% of trials. To disentangle the latent decision parameters that lead to these RT differences, we fit the LCA model to the movement RT and accuracy data, split into “fast” and “slow” trials, separately for each participant. We let accumulation rate and non-decision time vary between fast and slow trials; all other parameters (threshold, noise, leakage, and inhibition) were fixed between the two conditions, yielding 8 parameters in total^1^. The non-decision time differed significantly between fast and slow trials (92 ms vs. 245 ms, p < .001), indicating that in slow trials, the RTs incorporate additional latencies prior to initiation of the accumulation process and/ or between termination of the accumulation and motor initiation compared to fast trials. In addition, the rate of the accumulation was higher for fast than for slow trials (.944 vs. .900, p = .006). These results predict that, if attentional selection covaries with the motor decision process, (1) the increase of perceptual performance in fast trials should start earlier than in slow trials (due to the difference in the non-decision time), and (2) the rate of the increase should be slightly steeper in fast compared to slow trials (due to the difference in accumulation rate). Figure 3A shows the time course of the discrimination rate for DTs at the instructed target location time-locked to final cue onset, separately for fast and slow trials. First, perceptual discrimination performance in fast and slow trials did indeed differ in the expected way: shortly (∼30 ms) after final cue onset, discrimination performance was superior in fast compared to slow trials (p < .007). To quantify the temporal offset and the rise of the curves, we fit sigmoid functions to the relationship between time-smoothed discrimination rate and time relative to final cue onset (t_cue_; in the window from -100 to 350 ms), separately for fast and slow trials:

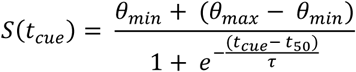

**Figure 3.**
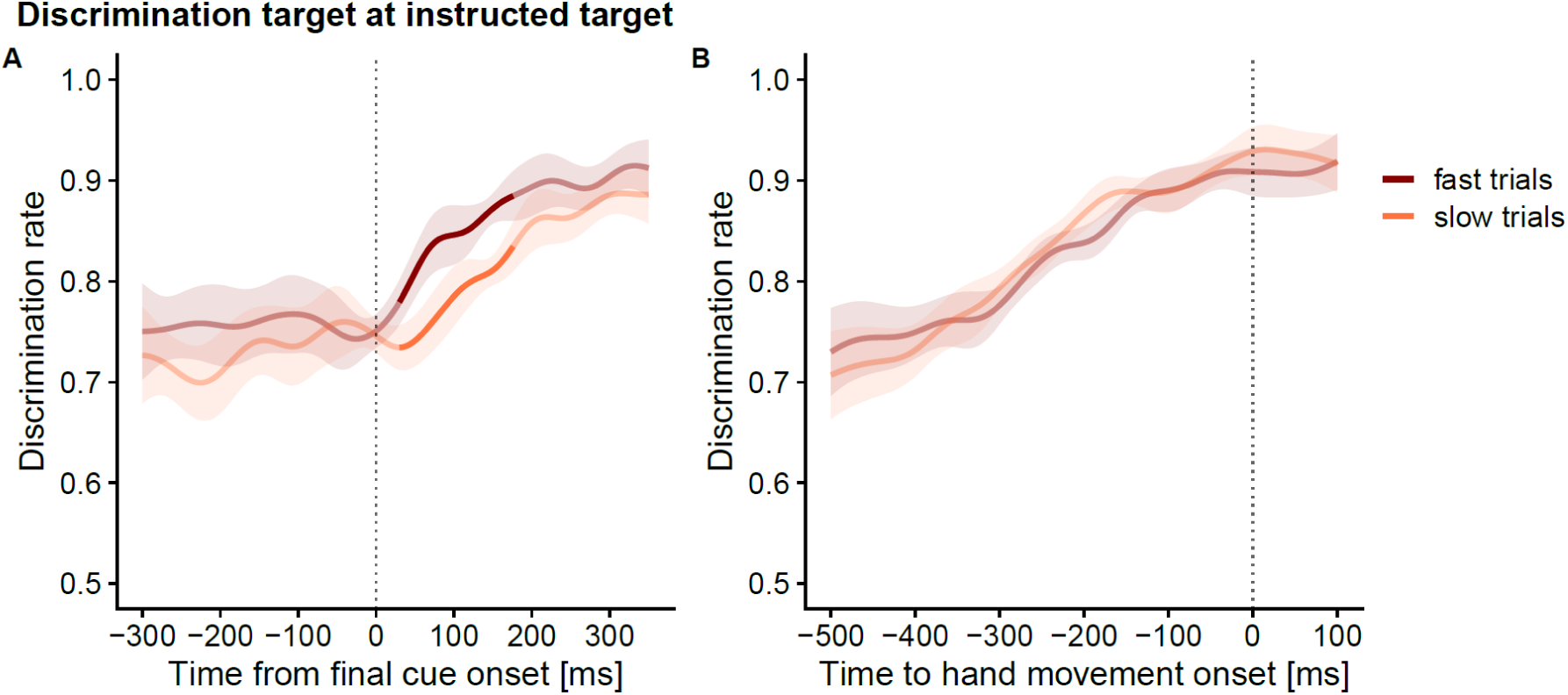
Time-course of visuospatial attentional in Experiment 1 for trials in which the DT was presented at the instructed target location, separately for fast (brown) and slowly initiated (orange) reaches. (**A**) shows the data time-locked to final cue onset and (**B**) time-locked to reach onset. Data are smoothed and weighted means (see Method section). Saturated segments denote time periods with significant differences (p < .05, weighted t-tests) between fast and slow trials. Shaded areas indicate the 95% confidence intervals.

The sigmoid functions contained four free parameters: a slope parameter τ that indicates the timescale over which the gradual change in perceptual discrimination occurred, a latency parameter t_50_ that indicates a shift of the sigmoid along the time axis, and two parameters θ_min_ and θ_max_, that denote the lower and upper limit of the sigmoid, respectively. Parameter estimates of the sigmoids showed that the gradual increase in discrimination performance started earlier in fast trials (t_50_ = 69 ms, 95% CI [61,77]) than in slow trials (t_50_ = 172 ms, 95% CI [151,193]), mirroring the large differences in non-decision time obtained from the LCA model. The increase also occurred within a shorter timescale for fast trials (τ = 34, 95% CI [27,41]) than for slow trials (71, 95% CI[50,92]), corresponding to the slightly faster accumulation rate (see Figure A2 in appendix). Notably, when time-locked to reach onset (Figure 3B), we did not observe statistically significant differences between the curves for fast and slow trials, thus suggesting that fast and slow trials may mainly differ with respect to the onset, rather than the rate, of evidence accumulation.

Taken together, the results of Experiment 1 first show that visuospatial attention was simultaneously allocated to both pre-cued target locations, indicating that multiple relevant reach targets can be selected in parallel. Second, once a motor goal was instructed, attention gradually increased at the goal location, suggesting that the representation of the instructed oal is strengthened. Third, the time-course of attentional selection correlated with reach latency and differed between reaches initiated at slow and fast latencies, in line with the proposal that attentional-perceptual processing reflects decisional processes leading to the selection of motor goals.

### Experiment 2

In Experiment 2, our hypothesis underwent a further test. If the time-course of attentional selection reflects evidence accumulation, it should be susceptible to factors known to be pertinent to sensorimotor decision-making. We thus modified our task such that the two pre-cued targets became the final reach target with different probabilities. For each participant, one cue color (e.g., blue) indicated that the respective location would become the final reach target with 80% probability (frequent target), and the other color (e.g., green), accordingly, indicated a 20% probability for the pre-cued location to become the reach target (rare target).

We first verified that our probability manipulation had the desired biasing influence on participants’ behavior. As expected, participants initiated their reaches faster when the frequent target was instructed as the final target compared to when the rare target became the final target (327 ms vs. 378 ms, *p* < .001). In addition, participants more often erroneously reached towards a deselected frequent target than towards a deselected rare target (2.64% vs. 0.88%, *p* < .001). Within the framework of sequential sampling models, such a bias is readily explained as a shift of the starting point of evidence accumulation towards the preferred option, so that less evidence is required to reach the decision threshold, consequently leading to shorter RTs (and fewer errors) for that option (Forstmann et al., 2016; Ratcliff et al., 2016). Alternatively, lower response latencies in frequent target trials might emerge from a higher accumulation rate or lower response thresholds. To account for these options, we again fit the LCA model to response latencies and accuracy in the movement task. We let accumulation rate and response threshold vary between frequent and rare target trials and estimated separate starting points for frequent and rare targets. We fixed all other parameters (leakage, inhibition, non-decision time, noise) between conditions.

Figure 4A illustrates mean and exemplary evidence accumulation trajectories for trials in which the frequent target was instructed as the final target and Figure 4B shows trials in which the rare target was instructed. The estimated starting point associated with the frequent target was significantly closer to the response threshold than that for the rare target (0.090 vs. 0.049, p < .001). In contrast, model fitting suggested neither a difference in accumulation rates (.891 vs. .875, p = .282), nor in response thresholds (0.345 vs 0.337, p = .283) between conditions. Thus, our model attributes the RT differences between frequent and rare target trials to diverging starting points, reflecting a response bias in favor of the frequent target.

**Figure 4.**
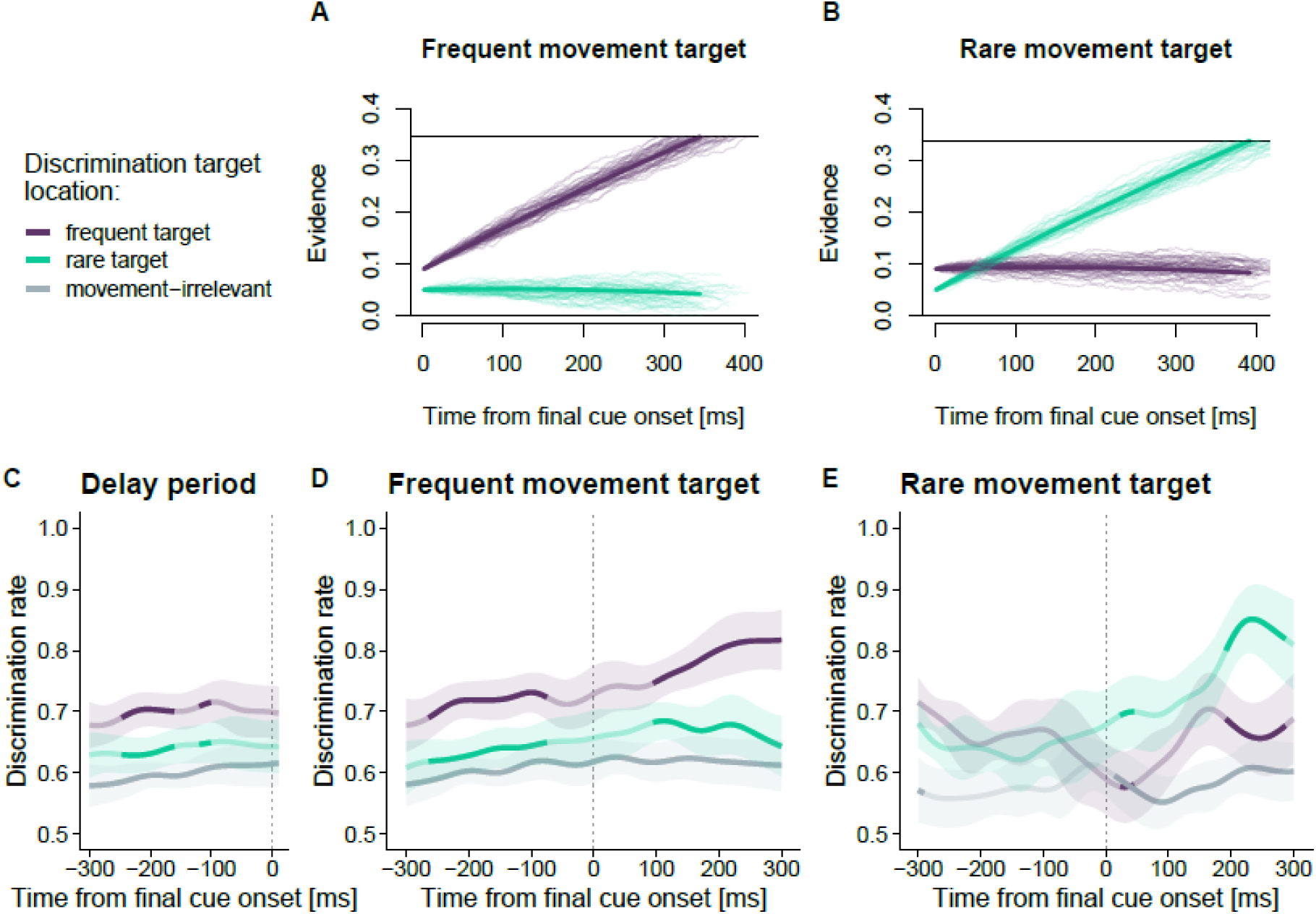
Time-course of visuospatial attentional in Experiment 2 for trials in which the DT was presented at the frequent target (purple), the rare target (green) and movement irrelevant locations (grey). (A) shows the delay period until specification of the final reach target, (**B**) shows trials in which the frequent target was instructed as reach target (80% of all trials in the experiment) and (**C**) trials in which the rare target was instructed (20% of all trials in the experiment). Data are smoothed and weighted means (see Methods section). Saturated segments denote time periods with significant differences (p < .05, weighted t-tests) between DT locations. Shaded areas indicate the 95% confidence intervals. **D** and **E** Mean (thick lines) and exemplary (thin lines) trajectories of the evidence accumulation. **D** Simulated trajectories for trials in which the frequent target (purple lines) became the instructed target. **E** Simulated trajectories for trials in which the rare target (teal lines) became the instructed target.

If the time-course of attentional allocation reflects the ongoing decision formation, such a bias should also be present in perceptual discrimination performance. Figure 4C shows the perceptual discrimination performance during the delay period, pooled across the frequent-target and rare-target conditions. In line with our hypothesis, spatial attention was already biased towards the frequent target during the delay period, evident in superior discrimination performance if the DT was shown at the frequent target compared to the rare target (∼ -250 to -160 ms; *p* = .028), and compared to movement irrelevant locations (*p* < .001) well before the final cue was presented. For trials in which the frequent target became the instructed reach target (Figure 4D), after final target onset discrimination performance further increased monotonically at the frequent target (now instructed as reach target) and was significantly higher than at the rare (now discarded) target (*p* < .001). Figure 4E shows the time-course of discrimination performance for trials in which the rare target became the finally instructed movement target. Note that due to the 80:20 distribution of reach target selection, the number of trials for this analysis is low by design. Therefore, data in this condition are inherently noisier compared to the frequent-target condition. Before final cue onset, perceptual discrimination performance was numerically higher at the frequent target compared to the rare target, though these differences were statistically not significant. Discrimination performance during the delay was, however, better at the frequent target compared to movement irrelevant locations (*p* = .013), while the differences between the rare target and the irrelevant locations failed to reach significance (p = .581), providing further evidence that the frequent target was preferentially processed during the delay. After final cue onset, discrimination performance increased at the rare, now instructed, target and was superior to both movement irrelevant locations (at ∼15 ms; *p* < .001) and to the frequent, now discarded, target (at ∼190 ms; *p* = .031).

These results indicate that our target probability manipulation modulated not only response latencies in the movement task, but also the time-course of visuospatial attention. During the delay, attention was biased towards the more frequent target and eventually (i.e., after final cue onset) further increased at the instructed target while the level of attention at the discarded target remained similar in this case. In case the rare target was instructed as the final reach target, this increase entailed a reversal of attentional priority of rare and frequent targets.

### Interim summary

So far, we have presented converging behavioral evidence that is consistent with the idea that perceptual discrimination performance, a marker for visuospatial attention, constitutes a time-extended marker of evidence accumulation during sensorimotor decisions, as formalized in sequential sampling models. Of note, while we modeled the accumulation process based on RT and accuracy data, the time-course of the simulated accumulation process shares at least three important characteristics with the experimentally obtained time-course of perceptual discrimination performance (Gold and Shadlen, 2007; O’Connell et al., 2018): First, perceptual discrimination at a specific location is best at the time of reach onset towards this location, which can be interpreted as the time point at which the decision is determined, and is analogous to the threshold that needs to be reached to trigger the associated response in the LCA model. Second, lower reach latencies were associated with a steeper and predominantly earlier increase of attentional enhancement at the instructed reach target. This observation is in line with an LCA model fit that located the parameter differences between trials with fast and slow reaction times mainly in the non-decision time, with only a small difference in the accumulation rate. Third, we observed that discrimination performance was susceptible to our probability manipulation with an initial bias towards the more frequent target, which over time became enhanced when movements had to be made to the frequent target and suppressed when movements had to be made to the rare target. In our model, this fact is well reflected in the different starting points for the frequent and rare targets (cf. Figure 4A vs. Figure 4D and Figure 4B vs. 4E).

As a caveat, the initial increase of perceptual discrimination performance after final cue onset at both the instructed and the discarded target location is not well captured by the model (see Figure 2A). Theoretically, certain parameter combinations could produce these results. For example, a slightly higher accumulation rate for the discarded target (but still lower than for the instructed target) and a larger mutual inhibition between response options could generate an initial increase in the discarded target as well, which is soon suppressed by inhibition stemming from the selected alternative. In an alternative view, it is not unlikely that those characteristics that could not be captured by the model reflect processes beyond the evidence accumulation for the motor decision. For example, to select the correct response, participants must first recall the respective locations of both color cues, and this retrieval from short-term memory may have driven the initial increase of perceptual performance (Herwig et al., 2010).

Finally, we acknowledge that the interpretation of our results as a gradual increase that correlates with evidence accumulation relies on data averaging, as the actual probed measurements are binary (correct/incorrect) responses. In principle, the attentional shift may be in fact a one-time, discrete step. Relatedly, the dominant view that neuronal activity in LIP resembles an evidence accumulation process has been challenged (Latimer et al., 2015). Rather, it has been suggested that the firing rate of LIP neurons undergoes rapid jumps that reflect discrete changes in the decisional state. However, (Shadlen et al., 2016) refuted this idea based on the observation that such discrete steps in activity are not observed when aligning the data to the end of the decision (i.e., movement initiation). Following this argument, if attention were shifted in a one-step process in response to a decision outcome, this should be evident in a step-like discontinuation of the gradual increase when aligning our data to reach onset, which we did not observe.

### Experiment 3

So far, our results suggest that visuospatial attention can be simultaneously deployed to multiple target locations. It is possible, however, that participants strategically only attend to one target in any given trial. Such a strategy would result in higher performance at the two pre-cued locations not because attention is concurrently distributed between them, but because, on average, in half of the trials, the choice of which particular location to attend would match the finally instructed movement target, resulting in higher performance when averaging across all performed trials. To reject this idea, it is necessary to confirm that both pre-cued locations are attentionally prioritized at the same time (Kramer and Hahn, 1995; Baldauf et al., 2006). In Experiment 3, we addressed this concern by replacing the discrimination task with a matching task (Kramer and Hahn, 1995). We presented DTs simultaneously at two locations, and participants reported whether identical or different DTs had been presented in the current trial. Thus, participants could only perform the task above chance level if they attend two spatial locations at the same time. We presented the two DTs either (1) at both pre-cued locations, (2) at one pre-cued and one movement-irrelevant location, or (3) at two movement irrelevant locations. If attention is split between the two potential targets, we expect matching performance to be superior if both DTs were presented at the pre-cued target locations compared to the other conditions.

Figure 5 shows the time-course of perceptual matching performance. Overall, performance was worse than in Experiments 1 and 2, indicative of higher task demands in the matching task. Furthermore, this finding has a simple mathematical reason: Assuming the discrimination rates are independent, the combined probability of obtaining a correct answer for both locations follows the multiplication rule for independent events, and is, thus, drastically smaller than for one correct answer. In Experiment 1 the discrimination rate during the delay period did not exceed 80%, which suggests a maximal matching rate of 64% for the two pre-cued locations. Nonetheless, during the delay, matching performance was better when both DTs were presented at the pre-cued locations compared to only one DT at a pre-cued location (at ∼ - 300 ms and ∼ -150 ms; *t*-tests: all *p* < .05, cluster non-significant) and compared to trials in which both DTs were displayed at movement-irrelevant locations (at ∼ - 70 ms; *t*-tests: all *p* < .05, cluster non-significant). After the final target had been specified, matching performance was superior when both DTs rather than just one (cluster *p* = .008) or none (cluster *p* = .015) occurred at pre-cued locations. Taken together, these results indicate that the two pre-cued target locations were simultaneously attentionally selected when the movement target was still uncertain and thus discard the possibility that the simultaneous attentional selection of both pre-cued targets in Experiment 1 is an artifact of trial averaging.

**Figure 5.**
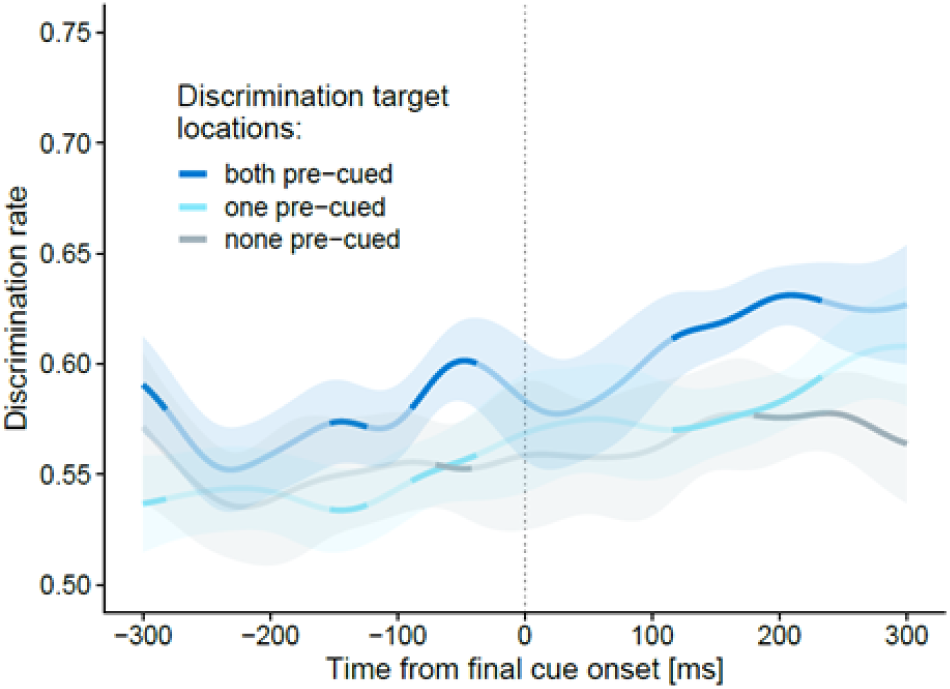
Time-course of discrimination accuracy in Experiment 3, separately for the conditions in which both DTs were shown at the pre-cued target locations (blue), in which only one DT was shown at a pre-cued target location (teal), and in which both DTs were shown at irrelevant locations (grey). Data are smoothed and weighted means (see Methods section). Saturated/bold segments denote time periods with significant differences (p < .05, weighted t-tests) between DT locations. Shaded areas indicate the 95% confidence intervals.

## Discussion

We examined the role of visuospatial attention during preparation of goal-directed movements in situations of target uncertainty. Participants performed center-out reaching movements to one of two pre-cued target locations while we probed perceptual discrimination performance at target and non-target locations at different time points during movement preparation.

We report four main results: First, attention was simultaneously allocated to both pre-cued targets, marked by higher discrimination performance at these than at other locations. Upon final goal designation, attention further increased monotonically at the goal location. Second, perceptual performance was highest around movement initiation, indicating a link between attention and motor preparation. Third, attention was sensitive to the probability of pre-cued locations to become the final goal, indicating that top-down information about the task structure biases attentional prioritization. Fourth, predictions from a sequential sampling model captured key temporal characteristics of attentional allocation across our experiments.

It is well established that visuospatial attention shifts towards goal locations prior to initiating saccadic eye or reaching movements (Baldauf and Deubel, 2010; Li et al., 2021). This phenomenon has been conceptualized to entail the formation of an attentional landscape or priority map that tags action-relevant locations through top-down weighting of visual input (Fecteau and Munoz, 2006; Ipata et al., 2009; Baldauf and Deubel, 2010). The present results suggest that the relationship between visuospatial attention and movement preparation goes beyond a link between a selected location and the respective movement to acquire it: visuospatial attention reflects the entire decision process that governs motor goal selection. We base this proposal on key similarities between the time courses of perceptual-attentional performance and evidence accumulation as formalized in sequential sampling models of decision-making (Bogacz, 2007; Gold and Shadlen, 2007; Ratcliff et al., 2016). Perceptual performance gradually increased at the goal location after the final target was specified, resembling evidence accumulation in favor of the motor goal. The nature of the gradual attentional increase depended on task- and response characteristics (i.e., the probability of a required answer and response latencies), as predicted by fits of the leaky competing accumulator model (Usher and McClelland, 2001) to the overt movement behavior. Finally, the peak of gradual increase coincided with the initiation of the movement, indicating a threshold in visual prioritization reflects a correlate of committing to a decision.

A link between visual-attentional performance and evidence accumulation has been previously proposed based on saccade-rather than reach planning (Jonikaitis et al., 2017). Given the intricate relationship between saccadic eye movements and visual perception for perceptual selection, this close link appears functionally useful, and is, indeed, deeply embedded neurally (Corbetta et al., 1998; Horwitz and Newsome, 1999; Nobre et al., 2000; Gold and Shadlen, 2000; Moore and Fallah, 2004; Gold and Shadlen, 2007; Ding and Gold, 2012). By contrast, reaching movements typically serve to manipulate the environment, have tactile consequences, are often not ballistic, and therefore require online control (Gallivan et al., 2018; Medendorp and Heed, 2019). Thus, although it is widely accepted that visuospatial attention supports motor preparation in hand movements (Allport, 1987; Land and Hayhoe, 2001; Wong et al., 2015), it is less straightforward to assume a direct and reciprocal relationship between visuospatial attention and manual actions than it is for saccades.

Just as LIP has a key role in oculomotor decisions, neurons in the parietal reach region (PRR) reflect decisions that are expressed by reaches (Andersen and Cui, 2009). Behavioral studies showed that attention can be allocated independently to saccade and reach targets, possibly indicative of separate, effector-specific networks for eye and hand movements that allocate attention (Jonikaitis and Deubel, 2011; Hanning et al., 2018). However, the degree to which the PPC comprises effector-specific subspaces remains debated (Medendorp and Heed, 2019). Recordings from LIP during a perceptual decision task revealed decision-related activity even when the decision was communicated via reaches rather than saccades (Lafuente et al., 2015), and inactivation of portions of LIP led to performance decrements in a free-choice reaching task (Christopoulos et al., 2018). Complementing these findings, neuroimaging work in humans has revealed a caudo-rostral gradient for eye versus limb movements with mostly common activation patterns for multiple effectors in posterior and differentiated activity in anterior PPC (Heed et al., 2011; Leoné et al., 2014). Thus, while PPC presumably entails effector-specific representations on the level of effector selection (Seegelke et al., 2021), effector-independent representations may underlie decisional processes related to motor goal selection as reflected in the time-course of visuospatial attention.

We observed that attentional prioritization was biased towards spatial locations that were likely to become the movement goal, and that reaches to these locations were initiated faster. Our computational model assigned the biasing influence of target frequency to the starting point of evidence accumulation, whereas other parameters of the decision, such as accumulation rate, were unaffected (see also Ratcliff et al., 1999; Leite and Ratcliff, 2011). Differences in starting points are usually explained in terms of increased neural activity that favors certain stimuli or actions prior to the decision process, such that less perceptual evidence is necessary to reach the decision threshold of the preferred option (Mulder et al., 2012; Forstmann et al., 2016). Neural activity in LIP correlates with factors that influence decision-making such as outcome probability (Platt and Glimcher, 1999; Yang and Shadlen, 2007), reward expectations (Kubanek and Snyder, 2015), and relative value (Sugrue et al., 2004), particularly during periods of response uncertainty early in trials. Thus, multiple biasing signals may be integrated in one common neural pool in which neural activity resembles a map of behavioral priorities that compete against each other for selection. Our present results suggest that visuospatial attention is tightly linked to this competition process. Notably, the discrimination target appeared at all locations with the same probability; accordingly, the presentation of the two relevant locations only concerned where participants had to reach but did not define where their visual discrimination would be probed. Thus, a position’s relevance for motor action overwrote bottom-up heuristics about potential discrimination target locations that participants could have used to prepare for the attention task (Druker and Anderson, 2010).

Finally, we observed superior perceptual performance when two concurrent discrimination targets appeared at the two pre-cued locations, arguing for the parallel encoding of potential motor goals (for similar results, see Baldauf et al., 2006). In strict interpretations of the concept of priority maps, multiple peaks on the map can signal multiple locations of interest. Nevertheless, it is usually assumed that one single option is chosen via a winner-takes-all mechanism (Hamker, 2005; Thompson and Bichot, 2005; Bisley and Goldberg, 2010). Consequently, attention should not be splitable in a sustained fashion. Indeed, in one study participants could equally split attention between two target locations (as measured by means of a memorization task), but only for a brief period of 100-150 ms. (Dubois et al., 2009). However, in other situations such as in our present task, it might be beneficial to maintain a stable representation of multiple locations (or even action plans, (Cisek, 2007; Cisek and Kalaska, 2010) to quickly initiate upcoming or correct ongoing movements (Nashed et al., 2014). This might be achieved by generating a sustained signal that is continuously fed back to networks involved in visual attention (Perry and Fallah, 2017).

To summarize, we demonstrate that the preparation of goal-directed reaching movements encompasses simultaneous allocation of visuospatial attention to multiple action-relevant locations. The time course of attentional prioritization was closely related to motor behavior and sensitive to target probability. We propose that our results fit well into the framework of competitive processing between simultaneously represented motor goals. In this framework, the time course of visuospatial attention is tightly coupled to spatial motor decisions and, thus, reflects the progression of decisions about motor goal selection, hence constituting a link of perceptual and motor aspects in sensorimotor decision-making.

## Acknowledgments

This work was supported by grants of the Deutsche Forschungsgemeinschaft (DFG) to T.H. (HE 6368/5-1) and C.Se. (SE 3004/1-1). We thank Florian Grünendahl and Lukas Söhring for help with data acquisition and Dennis Pauer for help with programming of the experiments. Data and code for the present paper are available at the Open Science Framework website https://osf.io/rxfjm/.

## Author contributions

T.H. and C.Se. designed research; C.Sc. performed research; T.H. contributed unpublished reagents/analytic tools; C.Sc. and C.Se. analyzed data; C.Sc. wrote the first draft of the paper; T.H. and C.Se. edited the paper; C.Sc., T.H., and C.Se. wrote the paper.

## Footnotes

1 It is also possible that RT differences result from distinct starting points for fast and slow trials. However, differences in starting points are usually conceptualized to capture biases for certain answers, for example when responses are associated with distinct desirability or expectations (Ratcliff et al., 2016; Forstmann et al., 2016). To address the possibility that participants exhibited a bias for one answer over the other, we also fit a model with varying starting points. The model fit revealed that starting points in slow and fast trials did not differ (p = .845). In addition, we obtained a superior fit for the model without the starting point parameter, as indicated by the lower Bayesian information criterion (BIC) value, -9601 vs. -9518 (Schwarz, 1978).

