## Supplemental Material for "Allocation of visuospatial attention indexes evidence accumulation for reach decisions"

### Appendix

#### Experiment 1

*Table A1.*
Experiment 1 *p* values and time points of identified clusters for pairwise differences between the discrimination target locations. The *p* value is given by the percentile of the nonpermuted cluster strength in the distribution of all permuted cluster strengths.

| | Comparison | Time (ms) | Cluster *p* value |
|---|---|---|---|
| Relative to final cue onset | | | |
| | irrelevant - discarded | -300 - 300 | < .001 |
| | irrelevant – instructed | -300 - 300 | < .001 |
| | instructed – discarded | -185 - -114 | .112 |
| | | 33 - 300 | < .001 |
| | instructed fast – slow | 30 - 176 | .007 |
| Relative to hand movement onset | | | |
| | irrelevant - discarded | -500 - -425 | .076 |
| | | -392 - 146 | < .001 |
| | irrelevant – instructed | -500 - 200 | < .001 |
| | instructed – discarded | -424 - 643 | < .001 |

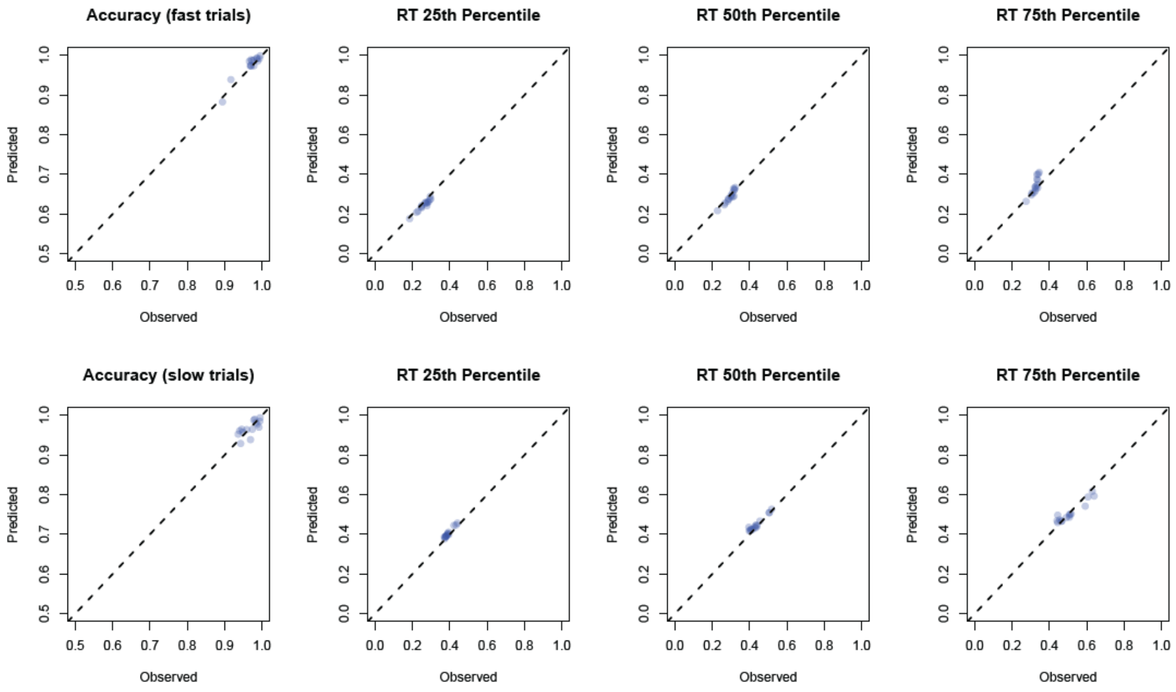


*Figure A1*. Illustration of the model fit for the Experiment 1 fast (top row) and slow (bottom row) trials as the relationship between empirical and predicted statistics (accuracy, 1st, 2nd, and 3rd RT quartile). Blue circles depict individual participant data and predictions.

*Table A2.*
Grid search starting values for the LCA model parameters (Experiment 1)

| | |
|---|---|
| Leakage | 0.3, 0.9 |
| Boundary | 0.2, 0.35, 0.5 |
| Noise constant | 0.2, 0.35, 0.5 |
| Inhibition | 0.3, 0.9 |
| Accumulation rate | |
| fast | 0.85, 0.9, 0.95 |
| slow | 0.85, 0.9, 0.95 |
| Nondecision time | |
| fast | 0.05, 0.15 |
| slow | 0.05, 0.15 |

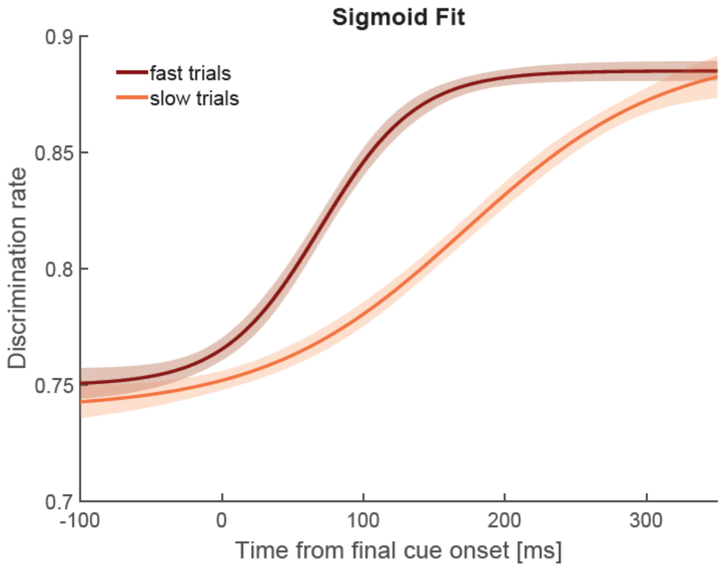


*Figure A2.* Sigmoid function fit to time-smoothed discrimination rate and time relative to final cue onset in the window from -100 to 350 ms, separately for fast and slow trials. The sigmoidal function contained four free parameters: a slope parameter $\tau$ for the timescale over which the gradual change in perceptual discrimination occurred, a latency parameter $t_{50}$ indicating a shift along the time axis, and $\theta_{min}$ and $\theta_{max}$, that denote the lower and upper limit of the sigmoid. Gradual increase in discrimination performance started earlier in fast trials ($t_{50}$ = 69 ms, 95% CI [61,77]) than in slow trials ($t_{50}$ = 172 ms, 95% CI [151,193]) and occurred within a shorter timescale for fast trials ($\tau$ = 34, 95% CI [27,41]) than for slow trials (71, 95% CI[50,92]).

#### Experiment 2

*Table A3.*
Experiment 2 *p* values and time points of identified clusters for pairwise differences between the discrimination target locations.

| | Comparison | Time (ms) | Cluster *p* value |
|---|---|---|---|
| During Delay | | | |
| | irrelevant - discarded | -300 - -247 | .112 |
| | | -178 - -122 | .111 |
| | irrelevant – instructed | -300 - 0 | < .001 |
| | instructed – discarded | -247 - -161 | .041 |
| | | -120 - -101 | < .331 |
| Frequent target | | | |
| | irrelevant - discarded | -35 - -26 | .555 |
| | | 44 - 161 | .006 |
| | | 188 - 258 | .119 |
| | irrelevant – instructed | -300 - 300 | < .001 |
| | instructed – discarded | -424 - 643 | .001 |
| | | 95 - 300 | < .001 |
| Rare target | | | |
| | irrelevant - discarded | -300 - -233 | .099 |
| | | -222 - -84 | .013 |
| | | 104 - 207 | .018 |
| | irrelevant – instructed | -300 - -291 | .581 |
| | | 14 - 300 | < .001 |
| | irrelevant - discarded | 23 - 45 | .517 |
| | | 192 - 287 | .021 |

*Table A4.*
Grid search starting values for the LCA model parameters (Experiment 2)

| | |
|---|---|
| Leakage | 0.3, 0.9 |
| Nondecision time | 0.5, 0.15 |
| Noise constant | 0.2, 0.35, 0.5 |
| Inhibition | 0.3, 0.9 |
| Starting point | |
| frequent target | 0, 0.05, 0.1 |
| rare target | 0, 0.05, 0.1 |
| Accumulation rate | |
| frequent target | 0.85, 0.9, 0.95 |
| rare target | 0.85, 0.9, 0.95 |
| Boundary | |
| frequent target | 0.2, 0.35, 0.5 |
| rare target | 0.2, 0.35, 0.5 |

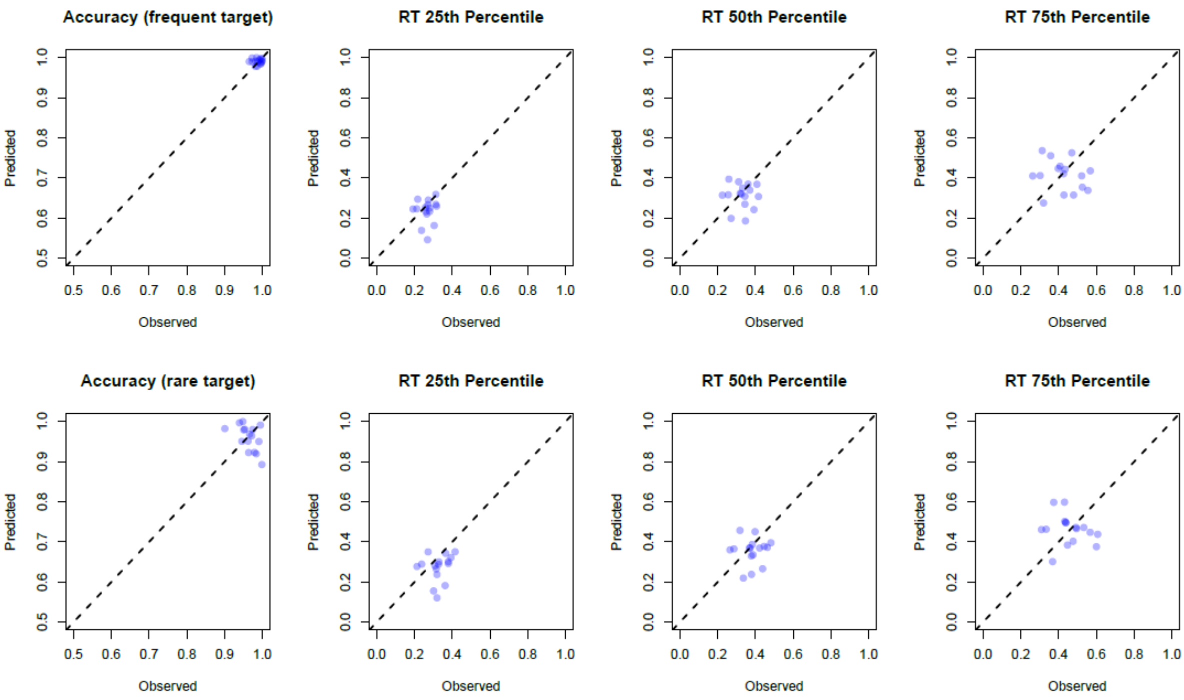


*Figure A3*. Illustration of the model fit for the Experiment 2 frequent (top row) and rare (bottom row) target trials as the relationship between empirical and predicted statistics (accuracy, 1st, 2nd, and 3rd RT quartile). Blue circles depict individual participant data and predictions.

#### Experiment 3

Table A5.
Experiment 3 *p* values and time points of identified clusters for pairwise differences between the discrimination target locations.

| | Comparison | Time (ms) | Cluster *p* value |
|---|---|---|---|
| Relative to final cue onset | | | |
| | both – none | -70 - -42 | .415 |
| | | 180 - 300 | .015 |
| | both – one | -300 - -282 | .477 |
| | | -153 - -124 | .412 |
| | | -89 - -38 | .200 |
| | | 116 - 233 | .008 |
